# Solving for X: evidence for sex-specific autism biomarkers across multiple transcriptomic studies

**DOI:** 10.1101/309518

**Authors:** Samuel C. Lee, Thomas P. Quinn, Jerry Lai, Sek Won Kong, Irva Hertz-Picciotto, Stephen J. Glatt, Tamsyn M. Crowley, Svetha Venkatesh, Thin Nguyen

**Author notes:** contributed equally (Samuel C. Lee, Thomas P. Quinn).

## Abstract

Autism spectrum disorder (ASD) is a markedly heterogeneous condition with a varied phenotypic presentation. Its high concordance among siblings, as well as its clear association with specific genetic disorders, both point to a strong genetic etiology. However, the molecular basis of ASD is still poorly understood, although recent studies point to the existence of sex-specific ASD pathophysiologies and biomarkers. Despite this, little is known about how exactly sex influences the gene expression signatures of ASD probands. In an effort to identify sex-dependent biomarkers (and characterise their function), we present an analysis of a single paired-end post-mortem brain RNA-Seq data set and a meta-analysis of six blood-based microarray data sets. Here, we identify several genes with sex-dependent dysregulation, and many more with sex-independent dysregulation. Moreover, through pathway analysis, we find that these sex-independent biomarkers have substantially different biological roles than the sex-dependent biomarkers, and that some of these pathways are ubiquitously dysregulated in both post-mortem brain and blood. We conclude by synthesizing the discovered biomarker profiles with the extant literature, by highlighting the advantage of studying sex-specific dysregulation directly, and by making a call for new transcriptomic data that comprise large female cohorts.

## 1 Introduction

Autism Spectrum Disorder (ASD) is a markedly heterogeneous condition with a varied phenotypic presentation and a spectrum of disability for those affected. As a neurodevelopmental disorder, the ASD syndrome is characterised by social abnormalities, language abnormalities, and stereotyped behavioural patterns Bailey et al. (1996). The presence of a genetic link in ASD etiology is well-established Miles (2011); Miyauchi and Voineagu (2013), first evidenced by ASD concordance among siblings and by a clear association between ASD and specific genetic disorders (e.g., Fragile X mental retardation) Bailey et al. (1996). This link has prompted a number of transcriptomic studies (e.g., Hertz-Picciotto et al. (2006); Glatt et al. (2012); Gupta et al. (2014)) to identify gene expression signatures (i.e., as a kind of biomarker) that might help elucidate the etiology of ASD and aid in its diagnosis (an important objective since early diagnosis and therapy is shown to improve outcomes in ASD Elder et al. (2017)). However, despite the number of transcriptomic studies performed, the pathophysiology and biomarker profile of ASD are still not known. Rather, these studies have tended to produce inconsistent results, suggesting wide heterogeneity among both the individual patients and the study populations. Indeed, ASD may not have one signature at all, but instead multiple diverging signatures Tylee et al. (2017a).

Transcriptomic studies of ASD probands typically use cells collected from either post-mortem brains or blood in order to estimate the mRNA abundance for thousands of gene transcripts (by way of microarray technology or massively parallel high-throughput sequencing (RNA-Seq)). Since many expressed transcripts are a precursor to structural or functional proteins, these studies can provide an insight into the functional state of a cell, capturing the common pathway for hereditary predisposition and environmental exposure. Although post-mortem brain studies have an advantage in that they look directly at the tissue of interest, blood-based studies can identify clinically useful biomarkers while also serving as a reliable proxy for gene expression in the brain Tylee et al. (2013) (though a complete understanding of ASD pathophysiology and its biomarker profile will likely require careful consideration of both lines of evidence). To date, more than a dozen studies have measured the transcriptomic profiles of ASD probands (and controls), the results of which have been summarised by two separate meta-analyses Ch’ng et al. (2015); Ning et al. (2015) and one “mega-analysis” Tylee et al. (2017a).

Sex is often called a risk factor for ASD, and it is stated that the risk for a male to have ASD is four to five times higher than that for females Werling et al. (2016); Christensen et al. (2016) (although the magnitude of this difference may be partly due to diagnostic biases Lai et al. (2015)). A similar observation, that the increased male risk is even higher among high-functioning ASD probands Fombonne (1999), likewise suggests that sex-specific mechanisms could influence ASD pathophysiology and its biomarker profile. Further evidence for sex-specific mechanisms is found in recent transcriptomic and functional-imaging studies. For example, Tylee et al., using transformed lymphoblastoid cell lines, found evidence for sex-specific differential regulation of genes and pathways among ASD probands Tylee et al. (2017). Similarly, Trabzuni et al. found sex-specific differences in alternative splicing in adult human brains, including for a well-known ASD risk gene NRXN3 Trabzuni et al. (2013). Functional brain connectivity studies using fRMI imaging have also identified sexual heterogeneity among ASD probands, showing dysregulation in sexually dimorphic brain regions across two large studies Floris et al. (2018); Lai et al. (2013). Moreover, recent work by Mitra et al. found evidence for pleiotropy between common single nucleotide polymorphisms (SNPs) for secondary sex characteristics and ASD risk, as well as sex heterogeneity on the X-chromosome, through a comprehensive SNP “mega-analysis” combining 12 individual data sets from diverse genetic backgrounds Mitra et al. (2016). Taken together, it seems plausible that sex could interact with other genetic and environmental factors to create sex-specific ASD pathophysiologies and biomarker profiles.

As ASD is more common in males, it suggests that females may have some underlying protection whereby a higher risk load is required for them to become afflicted Robinson et al. (2013). One hypothesis posits that ASD itself reflects a shift towards “extreme maleness” such that males are necessarily predisposed Baron-Cohen (2002). In support of this, females with ASD do harbour more (and larger) copy number variants than males with ASDs Levy et al. (2011), and moreover exhibit differential penetrance given the same genetic etiology Lionel et al. (2014) (although Mitra et al. found no evidence for an increased SNP load in females Mitra et al. (2016)). Unfortunately, however, the increased prevalence of ASD in males has led to the exclusion of females from many transcriptomic studies (e.g., Hu et al. (2009); Sarachana et al. (2010); Alter et al. (2011)), making it difficult to understand the male skew in ASD prevalence. Indeed, individual studies are often underpowered to detect subtle sex-specific differences (if they contain female subjects at all). When female subjects are included, sex is typically modelled as a simple covariate rather than an interaction term (i.e. the ASD-sex interaction), meaning that only sex-independent (and not sex-dependent) biomarkers are discovered. When male ASD is contrasted with female ASD, it typically involves loosely comparing simple sex-specific differences (e.g., differential expression present in males but not females, and *vice versa*) in a statistically anticonservative manner. To our knowledge, no study has looked at whether gene expression signatures show a sex-autism interaction across multiple studies and human tissues.

Using a single paired-end post-mortem brain RNA-Seq data set and a meta-analysis of six blood-based microarray data sets, we present an analysis of transcriptomic data that focuses on comparing sex-dependent and sex-independent ASD biomarkers (and the functional profiles thereof) across multiple tissues. By modelling the interaction of sex and ASD directly, we identify biomarkers (as well as functional pathways) that show sex-differences in ASD probands that are different than those in control subjects. Then, for those biomarkers that show no interaction, we pool male and female probands for a secondary sex-independent analysis. Our results suggest that, despite low power, some genes have FDR-adjusted significant sex-dependent interactions, while even more have significant sex-independent main effects. Subsequent pathway analysis further shows that these sex-independent biomarkers have substantially different biological roles than the sex-dependent biomarkers, and that some of these pathways are ubiquitously dysregulated in both post-mortem brain and blood.

## 2 Methods

### 2.1 Data acquisition

#### 2.1.1 RNA-Seq data

We searched for relevant publicly available RNA-Seq data using the Gene Expression Omnibus (GEO) Barrett and Edgar (2006) with the term (“expression profiling by high throughput sequencing”[DataSet Type] AND (“autism spectrum disorder”[MeSH Terms] OR “autistic disorder”[MeSH Terms])) AND “homo sapiens”[Organism] (query made January 2018). We restricted eligible data sets to those sequenced with paired-end and non-poly-A-selected libraries. After excluding any data sets that used cell lines or did not have female cases, only one experiment, GSE107241 Wright et al. (2017), remained. These data comprise a RiboZero Gold paired-end RNA-Seq data set from 52 postmortem dorsolateral prefrontal cortex tissue samples.

Prior to alignment and quantification, raw RNA-Seq reads were trimmed using Trimmomatic (docker image quay.io/biocontainers/trimmomatic:0.36-4) Bolger et al. (2014) and quality control metrics were recorded (before and after trimming) using FastQC (docker image biocontainers/fastqc:0.11.5) Andrews (2010). We aligned trimmed reads and quantified expression using Salmon (docker image combinelab/salmon:0.9.0) Patro et al. (2017) as run in pseudo-quantification mode with a k-mer index of length 31. For the reference, we concatenated a human coding reference (i.e., GRCh38.90.cds) with the corresponding non-coding reference (i.e., GRCh38.90.ncrna).

#### 2.1.2 Microarray data

We collected multiple microarray data sets to perform a meta-analysis of sex-autism interactions and main effects of ASD (i.e., sex-independent effects, where males and females are pooled). We referenced two prior meta-analyses Ch'ng et al. (2015); Ning et al. (2015), and one “mega-analysis” Tylee et al. (2017a), to prepare a list of data sets to study. Of these data sets, we excluded any study that (a) measured transcript expression from brain tissue, (b) had no female cases, (c) used cell lines (i.e., GSE37772 and GSE43076), or (d) treated cells with PPA (i.e., GSE32136). Six data sets remained after exclusion, as described in Table 1.

**Table 1:**
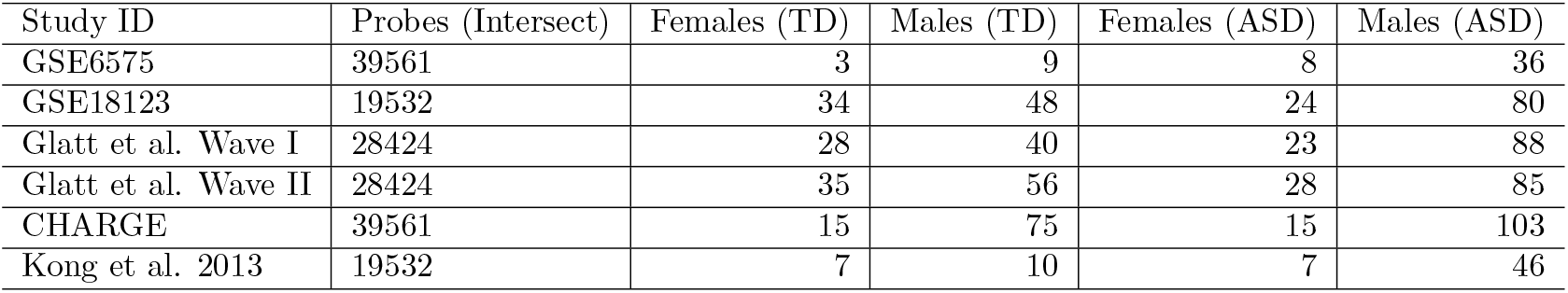
This table details all studies included in the meta-analysis, and the number of probes available after establishing a final probe set. All subjects with a labelled condition other than typically developed (TD) were assigned to the autism spectrum disorder (ASD) group, except for the two Glatt et al. data sets where “Type-1 errors” were assigned to the TD group.

| Study ID | Probes (Intersect) | Females (TD) | Males (TD) | Females (ASD) | Males (ASD) |
| --- | --- | --- | --- | --- | --- |
| GSE6575 | 39561 | 3 | 9 | 8 | 36 |
| GSE18123 | 19532 | 34 | 48 | 24 | 80 |
| Glatt et al. Wave I | 28424 | 28 | 40 | 23 | 88 |
| Glatt et al. Wave II | 28424 | 35 | 56 | 28 | 85 |
| CHARGE | 39561 | 15 | 75 | 15 | 103 |
| Kong et al. 2013 | 19532 | 7 | 10 | 7 | 46 |

Data acquired from the Gene Expression Omnibus (GEO) Barrett and Edgar (2006) (i.e., GSE6575 Gregg et al. (2008) and GSE18123 Kong et al. (2012)) were acquired already normalised and were not modified further. The other data sets (i.e., the Glatt et al. Wave I and Wave II data Glatt et al. (2012), the CHARGE study data Hertz-Picciotto et al. (2006), and the Kong et al. 2013 data Kong et al. (2013)) each underwent RMA normalization, quantile normalization, and base-2 logarithm transformation. All subjects with a labelled condition other than typically developed (TD) were assigned to the autism spectrum disorder (ASD) group, except for the two Glatt et al. data sets where “Type-1 errors” were assigned to the TD group. Note that, in crafting this dichotomy, some subjects assigned to the ASD group have delays that fall outside of the “spectrum” *per se*.

### 2.2 Differential expression analysis of RNA-Seq data

We used DESeq2 (Version 3.6) Love et al. (2014) to test for differential transcript expression within the Salmon-generated counts. We applied a conservative expression filter (i.e., at least 10 estimated counts per-gene in every sample) to the raw count matrix to ensure that the high variability of lowly expressed transcripts did not bias results due to the small group sizes. For each transcript that passed the expression filter, a model was fit using the formula ~ASD * Sex + Age (where Age is the age of death). Interaction and sex-independent main effects (i.e., of the ASD condition) were then extracted from the model by specifying the relevant contrasts to the DESeq2::results function. We corrected for multiple testing using the Benjamini-Hochberg procedure Benjamini and Hochberg (1995).

### 2.3 Meta-analysis of microarray data

Before proceeding with the meta-analysis, we established a set of probes (i.e., for each microarray platform) that represent genes also represented by probes in the other platforms. In other words, we established a final probe set based on the intersection of unique gene symbols present in all microarray platforms under study. Note that we resolved one-to-many mapping ambiguities by excluding any probe that mapped to multiple gene symbols.

For each microarray data set, and for each probe (i.e., of those representing genes found in all data sets), we performed differential expression analysis using limma (Version 3.34) Smyth (2004), applying the following steps: (1) fit a model with the formula ~ASD * Sex + Age where ASD and Sex are each two-level factors (except GSE6575, where the Age covariate is unknown), (2) define contrasts for the sex-autism interaction and for the sex-independent main effects (i.e., of the ASD condition), and (3) measure the differential expression for each contrast using the eBayes procedure.

Next, we transformed platform-specific probe p-values to HGNC symbol p-values using AnnotationDbi (available from Bioconductor Huber et al. (2015)). We resolved many-to-one mapping ambiguities by FDR-adjusting the minimum p-value of all probes for a given gene symbol (i.e., calculating a within-gene FDR correction). We then used Fisher’s method to perform a meta-analysis of the p-values obtained from the differential expression analysis. For *K* studies, Fisher’s method scores each gene based on (negative two times) the sum of the logarithm of the p-values:

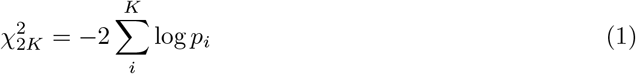

This score follows a χ^2^ distribution with 2*K* degrees of freedom Mosteller and Fisher (1948). Thus, for each gene, we computed a p-value directly from this score. We corrected for multiple testing using the Benjamini-Hochberg procedure Benjamini and Hochberg (1995).

### 2.4 Adjustment of latent batch effects

To ensure that latent batch effects did not inflate the discovery of false positives, we performed all analyses above with adjustment for batch effects using sva (Version 3.26) Leek et al. (2012); Leek (2014), applying the following steps: (1) estimate the number of surrogate variables while specifying the ASD * Sex interaction as the variable of interest and Age as an adjustment variable, (2) use the sva function (or, in the case of Salmon-generated counts, the svaseq function) to estimate the surrogate variables, and (3) include the surrogate variables in the differential expression model(s) described above. Generally speaking, using sva yielded more conservative results than not using sva. All tables and figures show results generated with sva except where otherwise noted.

### 2.5 Pathway analysis and knowledge integration

We performed pathway analysis using GSEA (Version 3.0) Subramanian et al. (2005a) in PreRanked mode with classic enrichment and 1,000 permutations. Enrichment scores were calculated for specific MSigDB (Version 6.1) Subramanian et al. (2005b); Liberzon et al. (2011) gene sets, including the curated KEGG (c2.cp.keggKanehisa et al. (2017)), Gene Ontology Biological Process (c5.bp) The Gene Ontology Consortium (2017), Reactome (c2.cp.reactome) Fabregat et al. (2018), and MSigDB Hallmark (h.all) Liberzon et al. (2015) sets.

Based on the nature of the analyses, input rank lists were prepared differently for the RNA-Seq and microarray results. For the RNA-Seq analysis, we ranked transcripts based on the p-value, *p*, and the magnitude of the fold-change, FC:

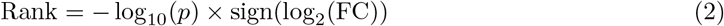

Then, these transcript-level ranks were converted into gene-level ranks based on the top transcript-level rank. For the microarray meta-analysis, we ranked genes using the χ^2^ test statistic (as calculated from Fisher’s method). Note that since this latter metric is agnostic to the direction of expression changes (i.e., only large χ^2^ test statistics suggest dysregulation), we focused here on pathways enriched with a positive score (effectively making this pathway enrichment test one-tailed).

## 3 Results

### 3.1 Evidence for sex-dependent autism biomarkers

By modelling the sex-autism interaction directly, we can detect gene expression signatures that have differential dysregulation in male ASD probands when compared with female ASD probands. In other words, we can find sexually dimorphic ASD biomarkers (e.g., a gene up-regulated in male ASD but not in female ASD, or *vice versa)*. Despite small study sizes (and disproportionately fewer females), we find some evidence for a sex-autism interaction among biomarkers, especially throughout the microarray meta-analysis data.

From the analysis of the RNA-Seq data derived from post-mortem brain tissue, we find no transcripts with significant (FDR-adjusted p-value < 0.05) sex-dependent dysregulation, although one of these transcripts showed a significant interaction prior to batch correction with sva. To illustrate what a sex-autism interaction might look like, Figure 1 shows the per-group expression profiles for the two transcripts with the largest interaction effect (i.e., those with the smallest corrected p-value). Table 2 characterises those transcripts with the most sex-dependent dysregulation.

**Figure 1:**
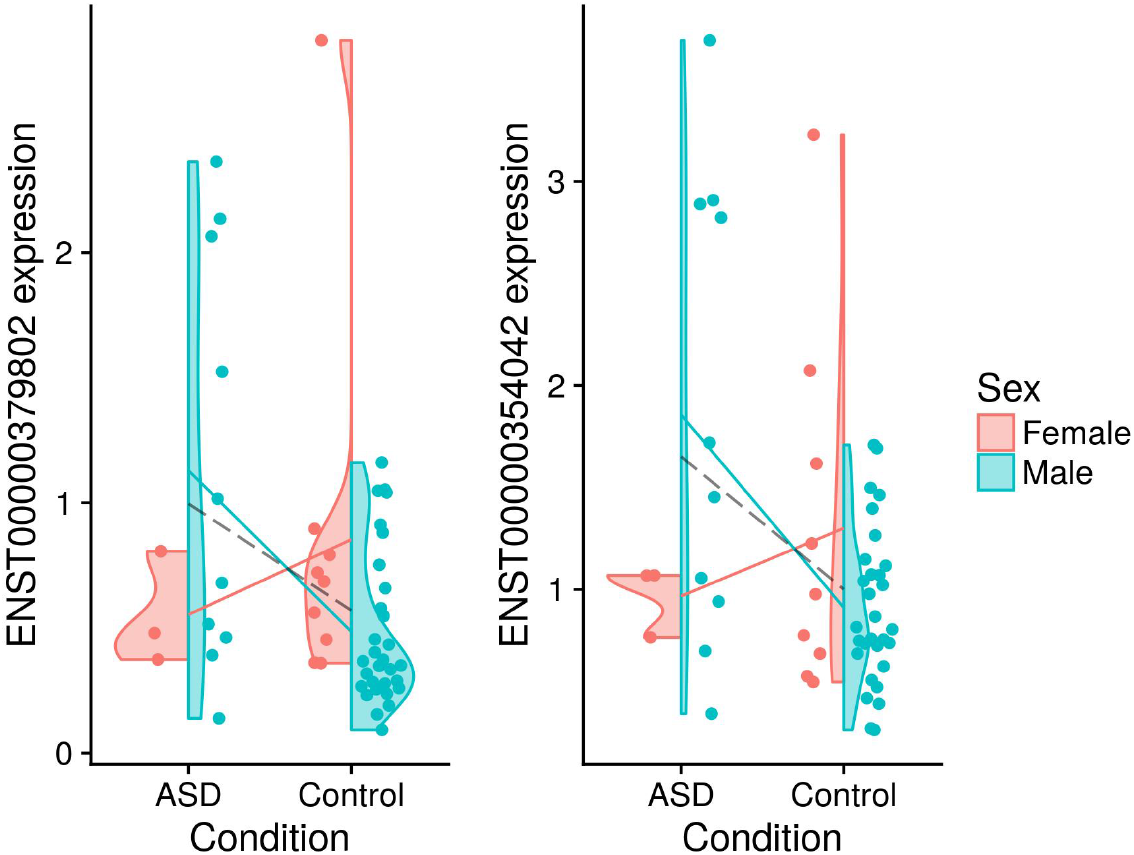
These violin plots show the base-2 logarithm-transformed expression for the two transcripts with the largest interaction effect from the RNA-Seq data (i.e., those with the smallest corrected p-value). The solid lines show sex-specific mean expression differences. The dashed line shows the sex-independent (i.e., pooled) mean expression difference.

**Table 2:** This table shows SVA-adjusted results for the sex-autism interaction for the RNA-Seq data (sorted by FDR-adjusted p-value). Note that FDR-adjusted p-values are also shown for an analysis performed without the adjustment of latent batch effects.

| Transcript ID | Gene symbol | Transcript biotype | Log 2 FC | P-adj SVA | P-adj (no SVA) |
| --- | --- | --- | --- | --- | --- |
| ENST00000354042 | SLC13A4 | protein_coding | 3.27 | 0.293 | 0.1136846 |
| ENST00000379802 | DSP | protein_coding | 3.19 | 0.293 | 0.6534814 |
| ENST00000262551 | OGN | protein_coding | 2.97 | 0.299 | 0.8169099 |
| ENST00000371625 | PTGDS | protein_coding | 1.74 | 0.299 | 0.0329544 |
| ENST00000223357 | AEBP1 | protein_coding | 1.85 | 0.529 | 0.8713166 |

From the meta-analysis of the blood-based microarray data, we find two genes with significant (FDR-adjusted) sex-dependent dysregulation: *TTF2* and *UTY*. Table 3 characterises those genes with the most sex-dependent dysregulation. Since for a meta-analysis by Fisher’s method, a large departure from the null (i.e., a very small p-value) in only one of several studies could cause the meta-analysis to post a significant result (i.e., even after FDR-adjustment) Tseng et al. (2012), it is useful to inspect visually how each study contributed to the results of the meta-analysis. For this, Figure 2 shows how each study contributed to the meta-analysis findings by plotting the aggregate Fisher score for each gene (of those with large sex-dependent dysregulation) along with the study-wise nominal significance (unadjusted p-value < 0.05). Notably, several of the most significantly dysregulated genes are at least nominally significant in more than one study.

**Table 3:** This table shows genes with the most sex-dependent dysregulation (and their chromosomal position), sorted by Fisher score and adjusted p-value. In addition, this table shows the Fisher score and adjusted p-value calculated for an analysis repeated without the adjustment of latent batch effects.

|  | Location | Fisher | Fisher p-adj | Fisher (no SVA) | Fisher p-adj (noSVA) |
| --- | --- | --- | --- | --- | --- |
| TTF2 | 1p13.1 | 52.16404 | 0.0105053 | 28.88686 | 1.0000000 |
| UTY | Yq11.221 | 50.59543 | 0.0198876 | 45.76688 | 0.1378710 |
| KCNJ8 | 12p12.1 | 48.17048 | 0.0528841 | 38.16932 | 1.0000000 |
| MAP1B | 5q13.2 | 48.15632 | 0.0531822 | 47.94878 | 0.0578051 |
| RAP2C | Xq26.2 | 47.70446 | 0.0637312 | 24.82099 | 1.0000000 |
| PAFAH1B1 | 17p13.3 | 45.45517 | 0.1559409 | 17.84249 | 1.0000000 |
| CRHR1 | 17q21.31 | 44.98624 | 0.1876599 | 43.46097 | 0.3416423 |
| NCS1 | 9q34.11 | 44.79693 | 0.2021903 | 30.46521 | 1.0000000 |
| CHST11 | 12q23.3 | 44.75342 | 0.2056750 | 21.19593 | 1.0000000 |
| SH3BGR | 21q22.2 | 44.61154 | 0.2174809 | 32.59585 | 1.0000000 |
| BNC2 | 9p22.3-p22.2 | 44.40363 | 0.2360031 | 39.81245 | 1.0000000 |
| RORA | 15q22.2 | 43.52113 | 0.3335702 | 34.11125 | 1.0000000 |
| HECA | 6q24.1 | 43.22311 | 0.3747481 | 33.12178 | 1.0000000 |
| FBRSL1 | 12q24.33 | 43.04625 | 0.4015007 | 35.53452 | 1.0000000 |
| PAK3 | Xq23 | 43.03339 | 0.4034965 | 43.20181 | 0.3780235 |
| ZC3H7B | 22q13.2 | 42.95711 | 0.4156536 | 35.03776 | 1.0000000 |
| CAMK1D | 10p13 | 42.80430 | 0.4411269 | 24.56439 | 1.0000000 |
| TMED10 | 14q24.3 | 42.55614 | 0.4858196 | 17.45529 | 1.0000000 |

**Figure 2:**
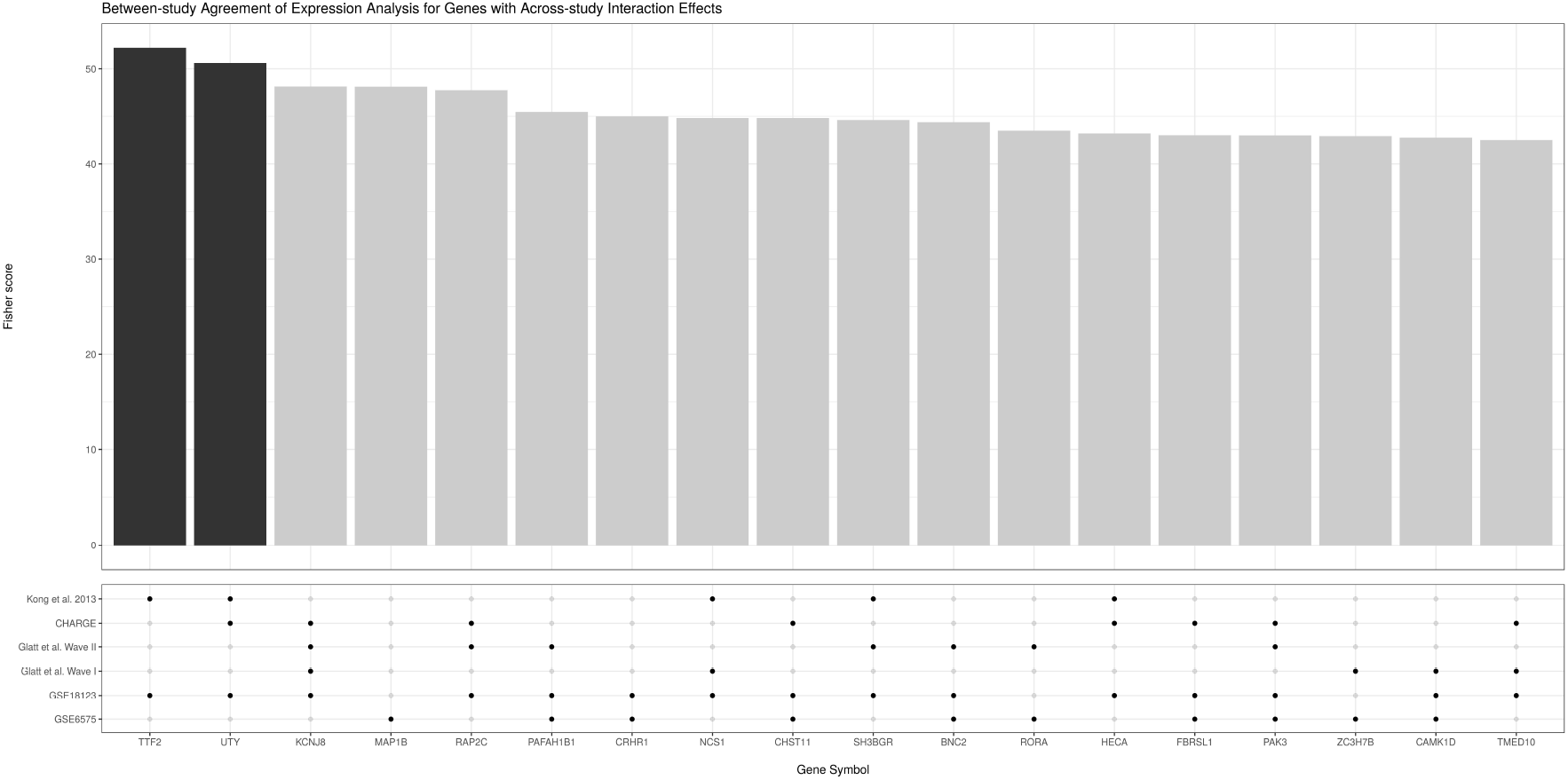
This figure shows the genes with the most significant sex-dependent dysregulation (i.e., a sex-autism interaction) according to the meta-analysis of the microarray data. Above, the bar plot shows the χ^2^ score for each gene as calculated using Fisher’s method (where the dark bars indicate that the gene has an FDR-adjusted p-value < 0.05). Below, the dot plot shows whether a gene showed a nominally significant sex-dependent dysregulation at an unadjusted p-value < 0.05 for a given study. Note that most genes selected for by the meta-analysis show at least nominal significance across multiple studies.

### 3.2 Evidence for sex-independent autism biomarkers

In situations where a sex-autism interaction is not detectable, we can proceed to measure main condition (i.e., sex-independent) effects by pooling male ASD probands with female ASD probands (and male controls with female controls), without having to model sex as a covariate. Genes with significant sex-independent main effects (i.e., of the ASD condition) have large unidirectional effect sizes in male ASD probands, female ASD probands, or both. Yet, because the interaction is tested first, we can interpret the main condition effects as sex-independent.

From the analysis of the RNA-Seq data derived from post-mortem brain tissue, we find seven transcripts with significant (FDR-adjusted p-value < 0.05) sex-independent differential expression. Of these, only one transcript showed significant up-regulation in ASD (with all others showing down-regulation). Figure 3 shows the expression profile for the two transcripts with the most significant sex-independent main effects (i.e., of the ASD condition). Table 4 characterises those transcripts with significant sex-independent dysregulation. Interestingly, several of the transcripts called differentially expressed by the analysis are annotated as non-coding RNA species.

**Figure 3:**
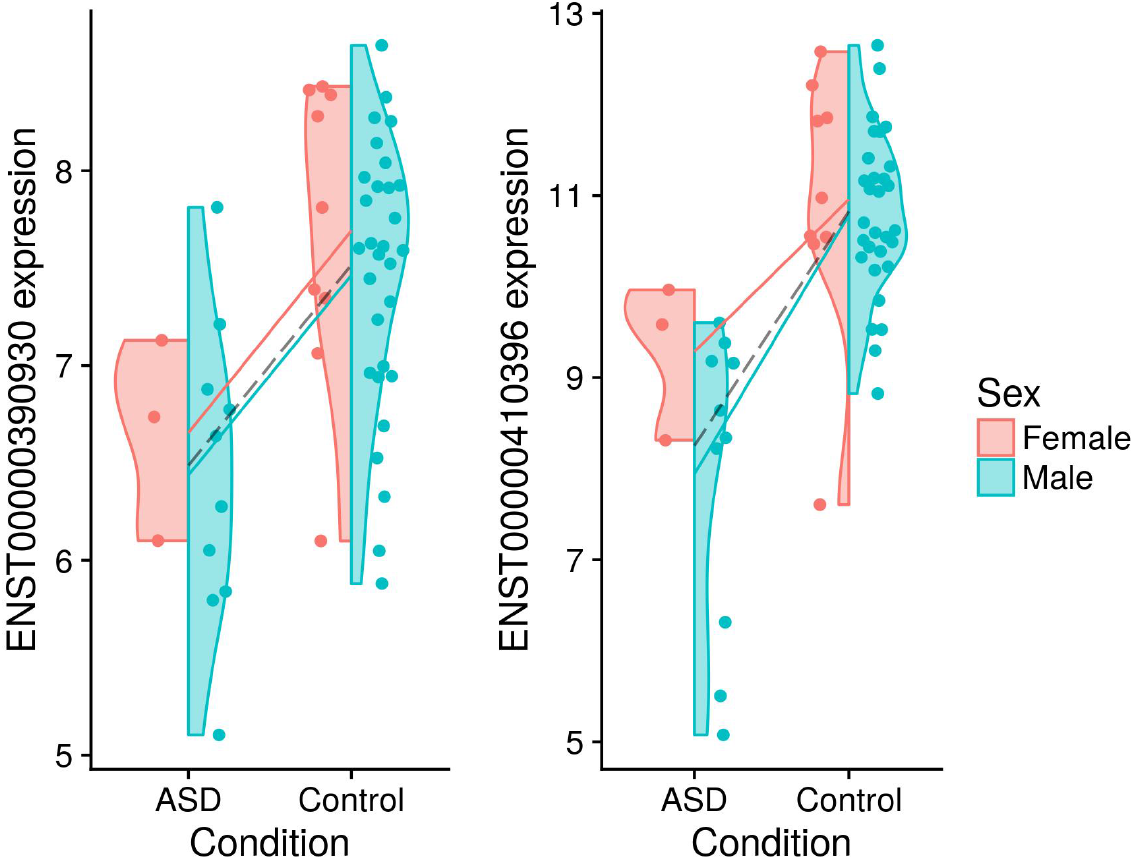
These violin plots show base-2 logarithm-transformed expression for the two most significant main effects (i.e., of the ASD condition) from the RNA-Seq data. The solid lines show sex-specific mean expression differences. The dashed line shows the sex-independent (i.e., pooled) mean expression difference.

**Table 4:**
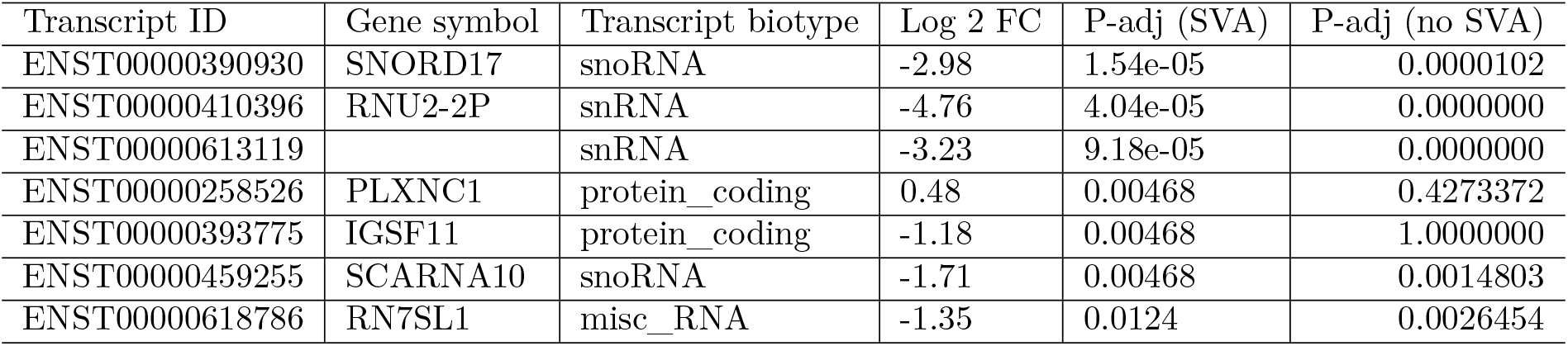
This table shows SVA-adjusted results for the main effects (i.e., of the ASD condition) for the RNA-Seq data (sorted by FDR-adjusted p-value). Note that FDR-adjusted p-values are also shown for an analysis performed without the adjustment of latent batch effects.

From the meta-analysis of blood-based microarray data, we find 21 genes with significant (FDR-adjusted) sex-independent dysregulation. Table 5 characterises those genes with the most sex-independent dysregulation. As in Figure 2, Figure 4 shows how each study contributed to the meta-analysis findings by plotting the aggregate Fisher score for each gene (i.e., of those with large sex-independent dysregulation) along with the study-wise nominal significance (unadjusted p-value < 0.05). Again, most genes selected as statistically significant by the meta-analysis are at least nominally significant in more than one study.

**Table 5:** This table shows genes with the most sex-independent dysregulation (and their chromosomal position), sorted by Fisher score and adjusted p-value. In addition, this table shows the Fisher score and adjusted p-value calculated for an analysis repeated without the adjustment of latent batch effects.

|  | Location | Fisher | Fisher p-adj | Fisher (no SVA) | Fisher p-adj (noSVA) |
| --- | --- | --- | --- | --- | --- |
| ARHGAP35 | 19q13.32 | 69.17663 | 0.0000083 | 59.97651 | 0.0004125 |
| GIMAP8 | 7q36.1 | 58.39735 | 0.0008000 | 52.71485 | 0.0083436 |
| UCHL3 | 13q22.2 | 55.85012 | 0.0023073 | 31.88589 | 1.0000000 |
| SPART | 13q13.3 | 53.75888 | 0.0054659 | 43.79029 | 0.2920570 |
| HNRNPA3P1 | 10q11.21 | 53.16493 | 0.0069742 | 54.55326 | 0.0039291 |
| ZNF721 | 4p16.3 | 53.02620 | 0.0073817 | 45.32102 | 0.1608751 |
| MAGED2 | Xp11.21 | 52.57098 | 0.0088931 | 31.43801 | 1.0000000 |
| UBE2A | Xq24 | 52.15816 | 0.0105264 | 24.84369 | 1.0000000 |
| KIF13B | 8p12 | 51.80723 | 0.0121459 | 44.99172 | 0.1830060 |
| POLR1A | 2p11.2 | 51.21815 | 0.0154371 | 35.12970 | 1.0000000 |
| PATL1 | 11q12.1 | 50.53892 | 0.0203385 | 37.55012 | 1.0000000 |
| COX19 | 7p22.3 | 50.48910 | 0.0207524 | 51.68452 | 0.0126954 |
| GNG5 | 1p22.3 | 50.42442 | 0.0213024 | 21.04799 | 1.0000000 |
| HNRNPF | 10q11.21 | 50.20526 | 0.0232786 | 52.19956 | 0.0102957 |
| MUM1 | 19p13.3 | 50.09134 | 0.0243757 | 38.59229 | 1.0000000 |
| MTERF4 | 2q37.3 | 49.77445 | 0.0277066 | 40.36576 | 1.0000000 |
| KLF1 | 19p13.13 | 49.50019 | 0.0309497 | 35.07655 | 1.0000000 |
| SART3 | 12q23.3 | 48.93549 | 0.0388576 | 51.89275 | 0.0116656 |
| EIF3A | 10q26.11 | 48.86929 | 0.0399046 | 48.66280 | 0.0429001 |
| ESF1 | 20p12.1 | 48.82351 | 0.0406442 | 40.26756 | 1.0000000 |
| TCEAL8 | Xq22.1 | 48.76924 | 0.0415389 | 30.32699 | 1.0000000 |
| RNF168 | 3q29 | 47.89156 | 0.0590766 | 40.39014 | 1.0000000 |
| NUCB2 | 11p15.1 | 47.52251 | 0.0684739 | 46.57846 | 0.0981743 |
| CCP110 | 16p12.3 | 47.21328 | 0.0774723 | 30.63996 | 1.0000000 |
| ZNF569 | 19q13.12 | 47.18319 | 0.0784042 | 35.01402 | 1.0000000 |
| CHP1 | 15q15.1 | 47.17381 | 0.0786939 | 46.71912 | 0.0928712 |
| ZC3H7B | 22q13.2 | 47.11959 | 0.0804103 | 31.75604 | 1.0000000 |
| GNPDA1 | 5q31.3 | 46.86648 | 0.0889439 | 39.70348 | 1.0000000 |
| ECI2 | 6p25.2 | 46.83204 | 0.0901676 | 54.27612 | 0.0044030 |
| VCP | 9p13.3 | 46.73363 | 0.0937667 | 33.68338 | 1.0000000 |
| ARHGAP8 | 22q13.31 | 46.70772 | 0.0947338 | 50.13461 | 0.0237714 |
| PGM1 | 1p31.3 | 46.58133 | 0.0996154 | 36.39139 | 1.0000000 |
| ZNF286A | 17p12 | 46.57586 | 0.0998268 | 31.41283 | 1.0000000 |

**Figure 4:**
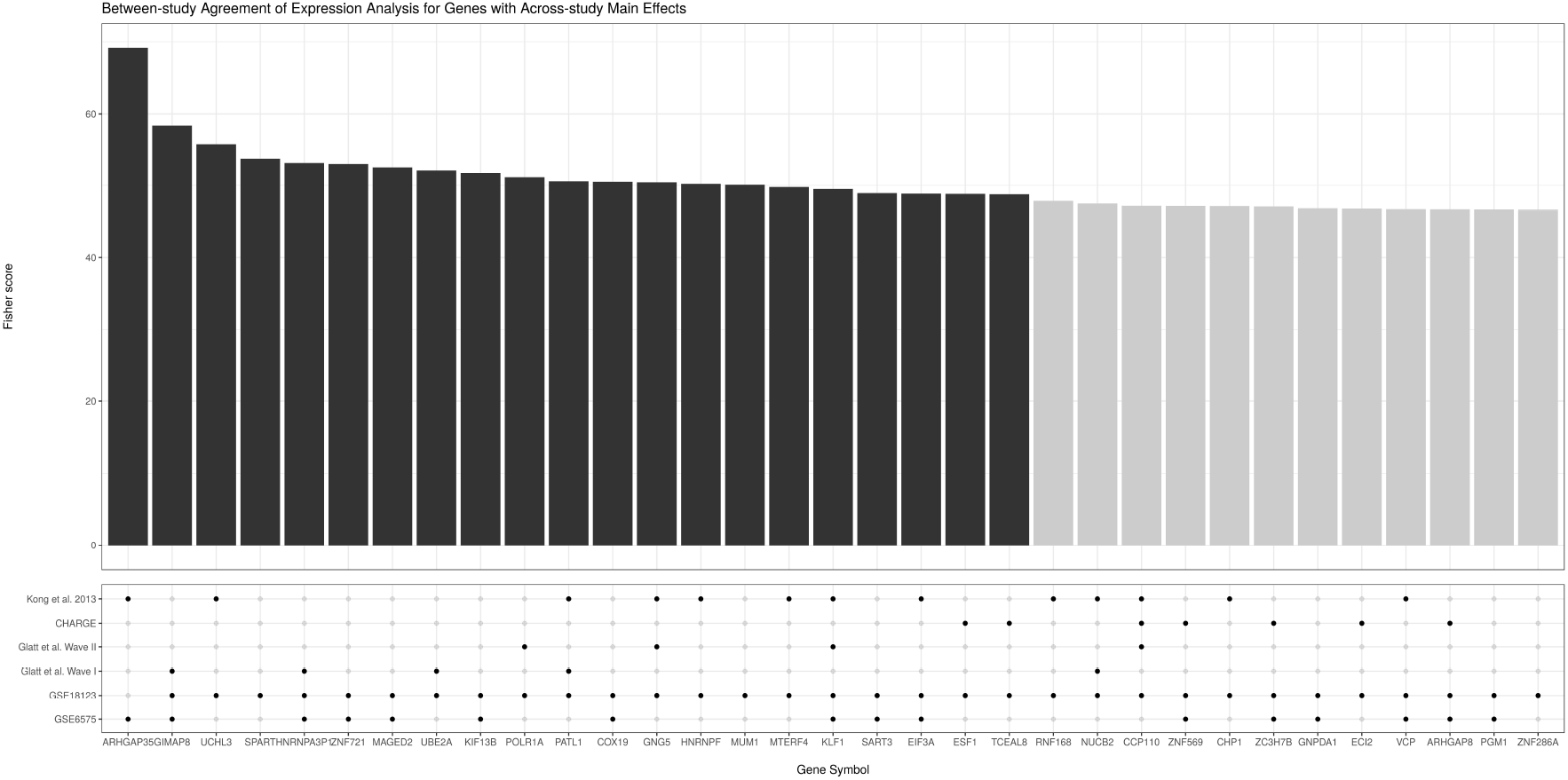
This figure shows the genes with the most significant sex-independent main effects (i.e., of the ASD condition) according to the meta-analysis of the microarray data. Above, the bar plot shows the χ^2^ score for each gene as calculated using Fisher’s method (where the dark bars indicate that the gene has an FDR-adjusted p-value < 0.05). Below, the dot plot shows whether a gene showed a nominally significant sex-independent main effect at an unadjusted p-value < 0.05 for a given study. Note that most genes selected for by the meta-analysis show at least nominal significance across multiple studies.

### 3.3 Pathway enrichment of ASD biomarkers

In an effort to summarise the biological relevance of the biomarker profiles generated above, we used the complete ranked lists of the differentially expressed transcripts (and genes) in four separate gene set enrichment analyses to identify common differentially regulated pathways. Four enrichment profiles were generated using the sex-dependent RNA-Seq (brain) biomakers, sex-independent RNA-Seq (brain) biomarkers, sex-dependent microarray (blood) biomarkers, and sex-independent microarray (blood) biomarkers.

Figure 5 shows the KEGG pathways enriched by the biomarkers as ranked by the analysis of the RNA-Seq data. For the sex-dependent biomarkers, nine pathways showed significant (FDR-adjusted p-value < 0.15) enrichment. For the sex-independent biomarkers, five pathways showed significant enrichment. Interestingly, all significant enrichment occurred in the same direction.

**Figure 5:**
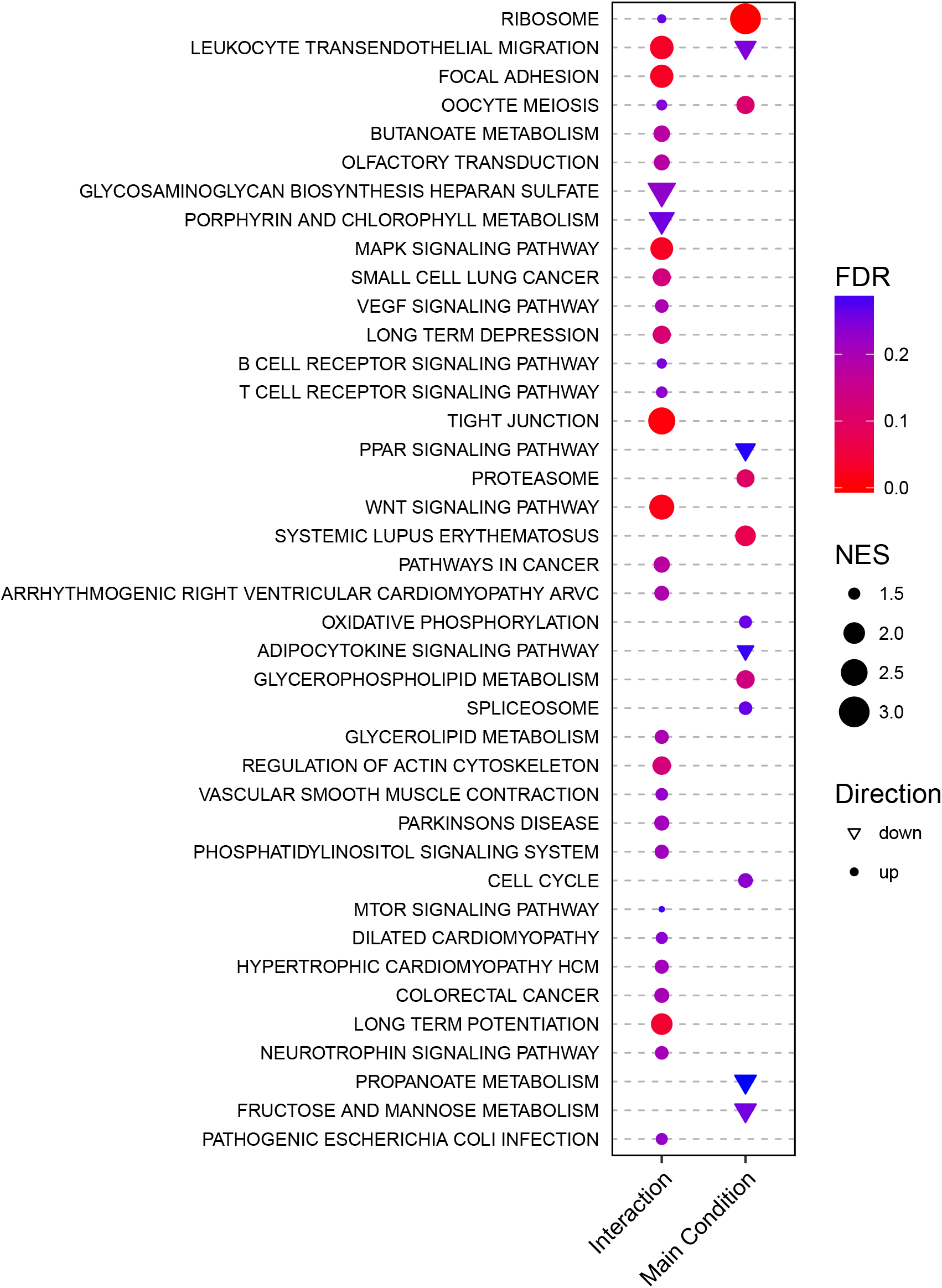
This dot plot shows results from a GSEA of the RNA-Seq data against the MSigDB KEGG pathways. For the two sets of results (i.e., the sex-autism interaction and the main effect), a KEGG pathway (y-axis) has a circle (or triangle) if it is enriched (or depleted). The size of the points indicates the absolute normalised enrichment score. The colour indicates the FDR. Note that only points with an FDR < 0.3 are plotted (see Methods).

Figure 6 shows the KEGG pathways enriched by the biomarkers as ranked by the analysis of the microarray data. For the sex-dependent biomarkers, one pathway (i.e., Alanine Aspartate and Glutamate Metabolism) showed significant (FDR-adjusted p-value < 0.30) enrichment. For the sex-independent biomarkers, thirty-six pathways showed significant enrichment. Note that because only positive (i.e., one-tailed) enrichments are considered for these data, an FDR-adjusted p-value < 0.30 is used here (see Methods for more details).

**Figure 6:**
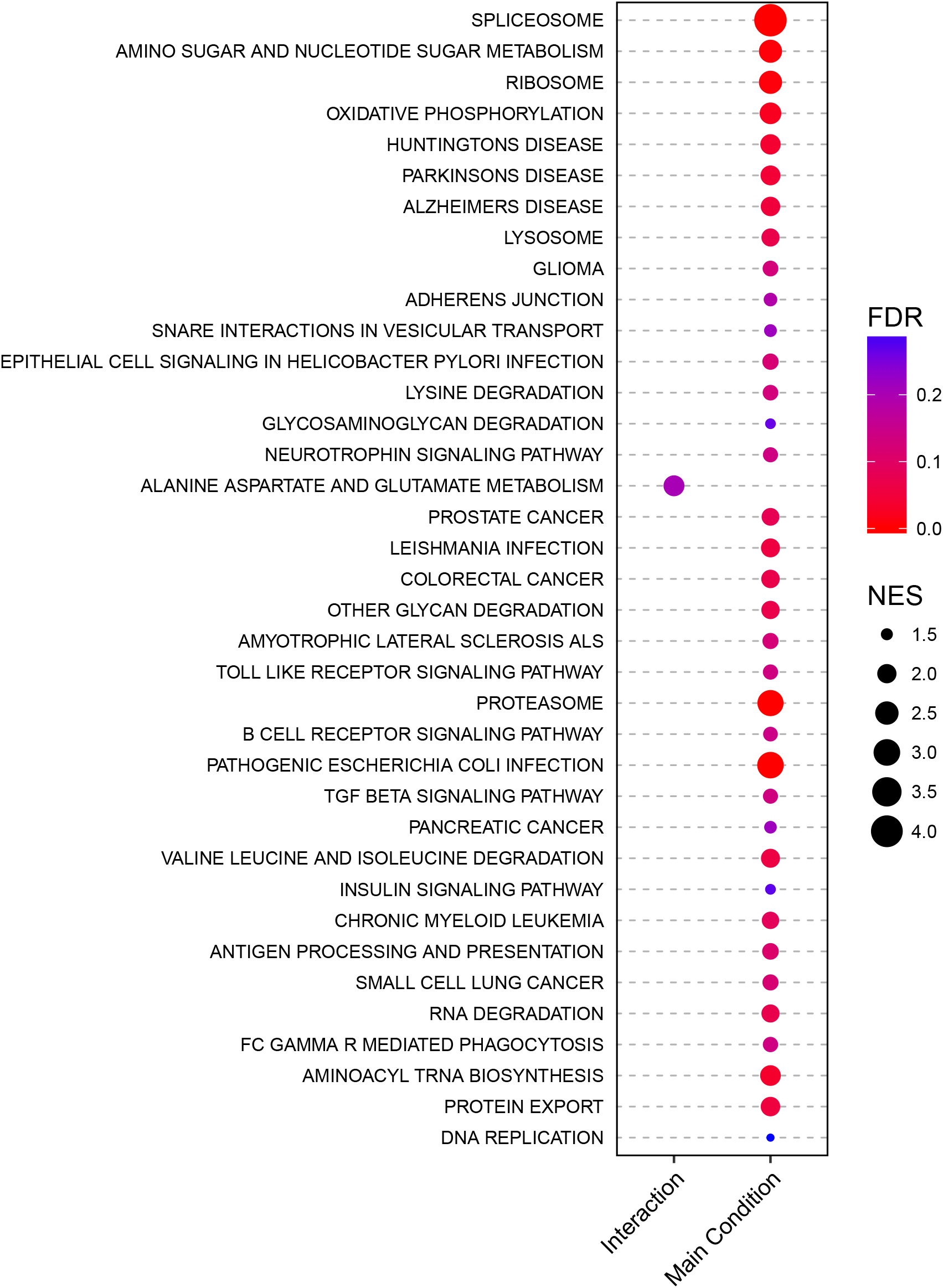
This dot plot shows results from a GSEA of the meta-analysis data against the MSigDB KEGG pathways. For the two sets of results (i.e., the sex-autism interaction and the main effect), a KEGG pathway (y-axis) has a circle if it is enriched. The size of the points indicates the absolute normalised enrichment score. The colour indicates the FDR. Note that only points with an FDR < 0.3 are plotted (see Methods).

Figure 7 compares the overlap between these significant pathways. For the sex-dependent analyses, no pathways are enriched in both the RNA-Seq and microarray data. However, for the sex-independent analyses, two pathways are enriched in both data. Interestingly, this agreement exists despite differences in the ranked lists, suggesting that ASD biomarker profiles may show some degree of higher-order conservation at the pathway-level that exists not only across multiple studies, but across multiple tissues (as well as multiple transcript quantification assays). Note that we also tested for enrichment among the Gene Ontology Biological Process, Reactome, and MSigDB Hallmarks gene sets, all of which show more examples of overlap between the separate sex-independent analyses (see the Supplementary Information for more details).

**Figure 7:**
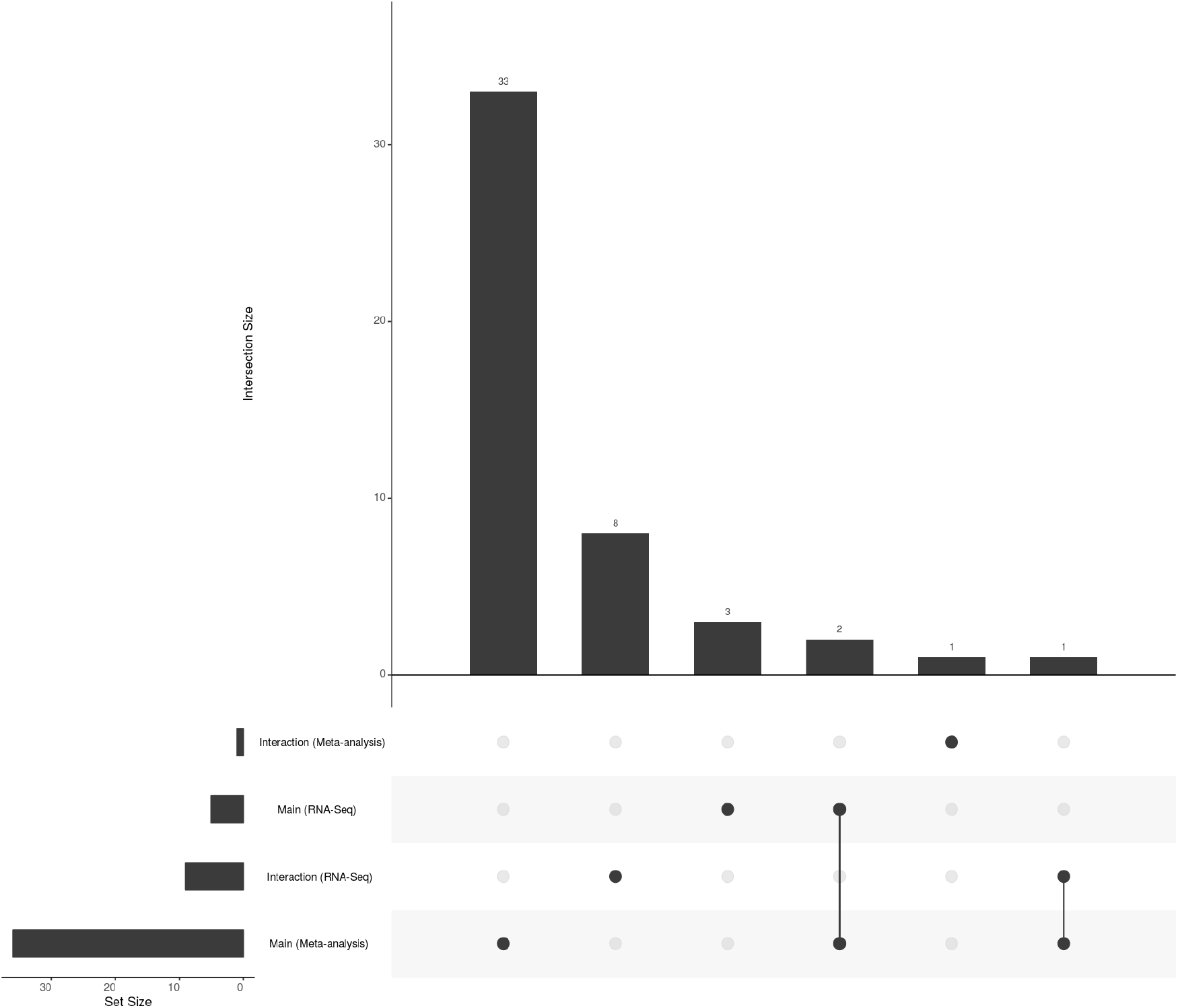
This UpSet plot Lex et al. (2014) shows set intersections (and their sizes) from a GSEA of four results against the MSigDB KEGG pathways. Set identity is indicated by the joined lines. Set size is indicated by the top bar chart. The bar chart on the left shows the total set size for each individual GSEA run. Results are filtered using a liberal FDR threshold of FDR < 0.15 for the RNA-Seq data and FDR < 0.3 for the meta-analysis data (see Methods).

## 4 Discussion

In this report, we present an analysis of several ASD transcriptomic studies, including an analysis of RNA-Seq data derived from post-mortem brain and a meta-analysis of six blood-based microarray data sets. Specifically, we focus on identifying both sex-dependent and sex-independent biomarker profiles for ASD by modelling the sex-autism interaction directly and secondarily measuring main effects of the ASD condition (i.e., sex-independent effects where males and females are pooled). In addition to identifying transcript (and gene) biomarkers, we use gene set enrichment analysis to summarise the observed dysregulation at the pathway level, contrasting sex-dependent pathway enrichment with sex-independent pathway enrichment. In doing so, we find evidence that ASD biomarker profiles may show some degree of higher-order conservation at the pathway level that exists not only across multiple studies, but across multiple tissues (and across multiple transcript quantification assays).

Despite small sample sizes in all studies, we found evidence for the existence of some sex-dependent biomarkers in human tissue. The meta-analysis identified two genes, *TTF*2 and *UTY*, with sexually dimorphic expression in the blood. One of these, *TTF*2, plays an important role in normal thyroid development De Felice and Di Lauro (2004). Interestingly, a loss of thyroid hormone homoeostasis has been linked to ASD Berbel et al. (2014); Khan et al. (2014). Since it is well-known that thyroid diseases have a sex-specific presentation Bauer et al. (2014), it seems plausible that thyroid abnormalities could contribute to a sexually dimorphic ASD signature. Some thyroid-disrupting environmental chemicals have also been linked to an altered risk for autism Lyall et al. (2017); Braun et al. (2014), including one study showing sexually dimorphic associations Lyall et al. (2017). The other, *UTY*, is a Y-chromosome gene (with considerable homology to an X-chromosome homolog), making any interpretation of its differential dysregulation difficult. Two other genes, *KCNJ*8 and *MAP1B,* had FDR-adjusted p-values very close to the pre-defined significance cutoff, warranting follow-up in another study. Although the RNA-Seq analysis did not yield any significant interactions, it is not surprising considering this data set contained only three female ASD probands. Nevertheless, the large (albeit non-significant) effect sizes warrant repeat studies with bigger cohorts and more female ASD probands.

By modelling the sex-autism interaction directly, we are able to follow-up the sex-dependent analysis with a secondary sex-independent analysis for any transcript (or gene) whose expression did not significantly interact with sex. In this scenario, we contrast the pooled male ASD probands and female ASD probands against the pooled male controls and female controls to calculate the main effects (which we can thus interpret as sex-independent biomarkers). Here, over twenty transcripts and genes exceeded the threshold for FDR-adjusted significance. Interestingly, for the RNA-Seq data, several of the significant biomarkers are not protein-coding genes (highlighting the value of using non-poly-A-selected libraries to quantify both coding and non-coding transcripts). For the microarray meta-analysis, several of the sex-independent biomarkers are associated with key neurodevelopmental processes, including some X-chromosome genes. For example, *MAGED*2, differentially expressed in ASD probands, is located on an X-linked intellectual disability hotspot (i.e., Xp11.2) Langnaese et al. (2001); Moey et al. (2016) (which, if causally relevant, could contribute to the male risk bias).

For both the RNA-Seq analysis and the microarray meta-analysis, we tested the ranked sex-dependent and sex-independent biomarker profiles separately for pathway-level enrichment. We found some pathway enrichment for the sex-dependent profiles, and even more for the sex-independent profiles. Importantly, very few of the enriched pathways were the same for both the interaction and main effects. This suggests that males and females exhibit unique pathway-level signatures that, if causally relevant, might further suggest the existence of both sex-specific and common ASD pathophysiologies. Although few KEGG pathways are enriched among the sex-dependent results, there are dozens of significantly enriched sex-dependent pathways across other tested gene sets (see Supplementary Information for more details). Among the sex-independent enriched pathways (for the meta-analysis results), there are a number of pathways for known neurodevelopmental and neurodegenerative diseases, including Huntingtons, Parkinsons, Alzheimers, and amyotrophic lateral sclerosis (ALS), suggesting that at least some of these ASD biomarkers may have functions important to general brain health. Considering that both unique and shared signatures (i.e., at the biomarker-level and pathway-level) exist among ASD probands, it seems plausible that molecular diagnostics could benefit from modelling sex-specific processes directly.

Although we found pathway enrichment to differ considerably between the sex-dependent and sex-independent biomarker profiles, we found that several sex-independent pathways (i.e., based on KEGG and other genes sets) were enriched across both the RNA-Seq and microarray data. Interestingly, this overlap exists despite the fact that analyses were performed on different human tissues (and with different transcript quantification assays). In fact, more than fifty Gene Ontology pathways were enriched among both sets of ranked sex-independent biomarkers (even though no gene products showed significant differential expression in both data). This overlap is consistent with a broad literature supporting common (and perhaps etiologically relevant) gene expression signatures across the widely heterogeneous population of ASD probands. If true, it seems plausible that molecular insights could further benefit from modelling pathway-level dysregulation directly (i.e., in addition to modelling conventional transcriptomic biomarkers).

When we compare our pathway enrichments to the previous ASD “mega-analysis” pathway enrichments Tylee et al. (2017b), we observe several complementary results. First, we found positive enrichment of the MAPK pathway in our sex-dependent RNA-Seq results, agreeing with the male-specific enrichment of Mek targets found in the Tylee et al. study Tylee et al. (2017b). Second, we found an enrichment of the ribosome-related pathway in both of our sex-independent analyses, agreeing with the ribosome-related pathway enrichment identified by the sex-independent “mega-analysis” Tylee et al. (2017b). Third, we found an enrichment of the Toll-like receptor (TLR) signalling pathway in our sex-independent meta-analysis results, agreeing with the TLR 3 and 4 signalling pathway enrichment identified by the sex-independent “mega-analysis” Tylee et al. (2017b). Importantly, these complementary results exist despite considerable differences in statistical methodology and data set inclusion.

Our analysis is not without limitations. First, although we used sva to adjust for latent batch effects, it is still possible that any number of remaining factors (or batch effects) could coincide with the diagnostic label (e.g., undocumented co-morbidities or medication use), thereby confounding the discovered biomarker profile. Second, as with any observational study, it is impossible to conclude whether the gene expression signatures (and their biological pathways) are causally related to ASD (or, likewise, the sex-autism interaction), rather than a result of the condition. Third, this analysis is likely under-powered to detect both sex-autism interactions and main effects, owing to the small sample sizes and disproportionately smaller female cohorts. Yet, based on the extant literature (which clearly highlights sex as an ASD risk factor) and the results published here, we believe that modelling the sex-autism interaction should become a mainstay of ASD transcriptomic research. Advantageously, as shown here, interaction modelling is compatible with the most commonly used softwares for batch-effect correction Leek et al. (2012), RNA-Seq analysis Love et al. (2014), and microarray analysis Smyth (2004). Yet, this analytical technique cannot offer any benefit if transcriptomic studies continue to systematically exclude female subjects (Hu et al. (2009); Sarachana et al. (2010); Alter et al. (2011)). Although there seems to exist a strong skew in the prevalence of male ASD, this very fact underlies the importance of studying female ASD: a complete understanding of the molecular basis of ASD will require the intentional study of both sex-dependent and sex-independent mechanisms, as well as their differences and commonalities.

## 5 Acknowledgements

SCL and TPQ designed the experiments, performed the analyses, and drafted the manuscript. JL provided statistical expertise and helped revise the manuscript. SWK, IHP, and SJG contributed data and helped revise the manuscript. TMC, SV, and TN supervised the project and helped revise the manuscript. This research was partially supported by the Australian Government through the Australian Research Council’s Linkage Projects funding scheme (LP140100240). SWK is supported by a grant from the National Institute of Health (R01MH107205). IHP is supported by grants from the National Institute of Environmental Health Sciences (1R01ES015359; 2R01ES015359; 1P30ES023513; 3P01ES011269), the National Institute of Health (UG3OD023365), and the United States Environmental Protection Agency (RD83543201).

